# Large-scale Genetic Characterization of a Model Sulfate-Reducing Bacterium

**DOI:** 10.1101/2021.01.13.426591

**Authors:** Valentine V. Trotter, Maxim Shatsky, Morgan N. Price, Thomas R. Juba, Grant M. Zane, Kara B. De León, Erica L. Majumder, Qin Gui, Rida Ali, Kelly M. Wetmore, Jennifer V. Kuehl, Adam P. Arkin, Judy D. Wall, Adam M. Deutschbauer, John-Marc Chandonia, Gareth P. Butland

## Abstract

Sulfate-reducing bacteria (SRB) are obligate anaerobes that can couple their growth to the reduction of sulfate. Despite the importance of SRB to global nutrient cycles and their damage to the petroleum industry, our molecular understanding of their physiology remains limited. To systematically provide new insights into SRB biology, we generated a randomly barcoded transposon mutant library in the model SRB *Desulfovibrio vulgaris* Hildenborough (DvH) and used this genome-wide resource to assay the importance of its genes under a range of metabolic and stress conditions. In addition to defining the essential gene set of DvH, we identified a conditional phenotype for 1,137 non-essential genes. Through examination of these conditional phenotypes, we were able to make a number of novel insights into our molecular understanding of DvH, including how this bacterium synthesizes vitamins. For example, we identified DVU0867 as an atypical L-aspartate decarboxylase required for the synthesis of pantothenic acid, provided the first experimental evidence that biotin synthesis in DvH occurs via a specialized acyl carrier protein and without methyl esters, and demonstrated that the uncharacterized dehydrogenase DVU0826:DVU0827 is necessary for the synthesis of pyridoxal phosphate. In addition, we used the mutant fitness data to identify genes involved in the assimilation of diverse nitrogen sources, and gained insights into the mechanism of inhibition of chlorate and molybdate. Our large-scale fitness dataset and RB-TnSeq mutant library are community-wide resources that can be used to generate further testable hypotheses into the gene functions of this environmentally and industrially important group of bacteria.

## INTRODUCTION

Sulfate-reducing bacteria (SRB) are present in diverse anoxic environments including the deep ocean, where they are responsible for a considerable fraction of carbon mineralization (Muyzer and Stams, 2008), and in the human gut, where they produce hydrogen sulfide (Kushkevych et al., 2020). Utilization of SRB has been extensively explored for bioremediation (eg. heavy metals, radionuclides) by metabolism-dependent mechanisms and/or bioaccumulation (Joo et al., 2015; Mikheenko et al., 2008; Rückert, 2016; Yong et al., 2002). In the oil and gas industry, the activity of SRB leads to undesirable effects including souring of oil and corrosion of pipelines (Kip and van Veen, 2015; Thrasher and Vance, 2005). Given their importance, it is imperative that we develop a detailed gene-level characterization of SRB to understand and control their activities in diverse environments.

Much of our molecular understanding of SRB is derived from studies in the model *Desulfovibrio vulgaris* Hildenborough (DvH), which was the first SRB to have its genome sequenced (Heidelberg et al., 2004). DvH has an established genetic toolkit including a markerless genetic exchange system (Keller et al., 2009), conceptual and predictive models of gene regulation and signal transduction (Rajeev et al., 2011; Turkarslan et al., 2017) and mapped networks of protein-protein interactions (Shatsky et al., 2016a, 2016b). Despite these advances, there remains considerable gaps in our understanding of DvH, and, hence, SRB as a whole.

Transposon mutagenesis is a powerful genetic tool for generating a large collection of mutant strains, and the measured phenotypes of these strains can be used to infer gene functions. In SRB, an ordered transposon library has been generated and characterized in *Desulfovibrio alaskensis* G20 (Kuehl et al., 2014). This collection was subsequently used to gain new insights into the electron transfer complexes of this bacterium (Meyer et al., 2014; Price et al., 2014). In addition, the transposon insertion sequencing approach (Tn-seq), whereby the abundance of thousands of mutants are assayed simultaneously through next-generation sequencing (van Opijnen and Camilli, 2013; van Opijnen et al., 2009), has been applied in the human-associated *Desulfovibrio piger* to characterize its metabolic niche (Rey et al., 2013) and in DvH to measure phenotypes under two conditions (Fels et al., 2013). In these previous Tn-seq studies, phenotypes were not detected for most genes in the respective genomes, partly because of the limited number of conditions assayed. More recently, the random barcode transposon-site sequencing (RB-TnSeq) approach has been developed which simplifies the measurement of mutant phenotypes across many conditions (Price et al., 2018; Wetmore et al., 2015), through the use of barcode sequencing or BarSeq (Smith et al., 2009).

In this work, we report the generation of an RB-TnSeq library in DvH and its use in generating a large gene-phenotype map using the BarSeq fitness assay across 757 experiments. These experiments represent 244 unique growth conditions including changes in respiratory and fermentative growth conditions, growth with different nitrogen and essential nutrient sources, and growth during exposure to various stressors. Through the investigation of this large gene-phenotype dataset in DvH, we define the essential gene set of this bacterium and derive specific new insights into its metabolism, regulation, and stress response.

## RESULTS

### DvH essential genes in the wild-type and Δ*upp* backgrounds

Genes with few or no transposon insertions are likely essential for viability under the growth conditions used to select the mutations. To identify the essential gene set of DvH, we first generated five transposon mutant libraries in the wild-type background on lactate-sulfate rich growth medium using a barcoded variant of the *Tn*5-RL27 transposon (henceforth called *Tn*5) (Larsen et al., 2002). Across all libraries, we generated 116 million Tn-seq reads with a *Tn*5 transposon insertion that mapped to the DvH genome, with the median gene represented by 17,823 reads. We used a previously described approach that estimates essential genes based on a number of criteria including gene length, insertion density, insertions in the central portion of the gene (as insertions near the 5’ and 3’ ends may not disrupt the gene’s function), and gene uniqueness (as we cannot discriminate insertions in highly repetitive regions) (Price et al., 2018; Rubin et al., 2015). In total, we identified 399 likely essential genes in the wild-type strain (Supplementary Table 1). Of these 399 genes, 322 (81%) have reduced transposon insertion coverage in a prior DvH transposon-sequencing experiment with the same growth medium (Fels et al., 2013).

We next constructed 24 *Tn*5 transposon mutant libraries in the JW710 strain background. Combined, we generated 154 million reads with a transposon and a mapped location of each in the DvH genome, with the median gene represented by 27,580 sequencing reads. JW710 contains a deletion of *upp* (DVU1025), encoding uracil phosphoribosyltransferase, a component of the pyrimidine salvage pathway. JW710 has been adopted as a commonly used base strain in which to perform counter-selection with resistance to 5-fluorouracil, a toxic pyrimidine (Keller et al., 2009). Using the same criteria applied to the wild-type transposon insertion data, we identified 436 likely essential genes in the JW710 background (Supplementary Table 1), of which 380 were in common with the wild-type background. The 380 genes that are shared between the two strains are a robust estimate of the essential gene complement of DvH and are enriched in general cellular processes such as protein synthesis and cell envelope functions (Figure 1A). To explore these genes further, we compared each to the Database of Essential Genes (DEG) (Zhang et al., 2004), which contains experimentally determined essential genes for dozens of diverse bacteria (all are non-SRB). We found that the majority of these 380 genes perform more general functions unrelated to sulfate reduction, as 271 have homologs that have been identified as essential in non-SRB. Only 109 of the DvH essential genes did not have a good homolog in DEG (Materials and Methods), and these include a number of well-known genes directly involved in sulfate reduction including *dsrAB* (dissimilatory sulfite reductase) and *sat* (sulfate adenylyltransferase).

**Figure 1.**
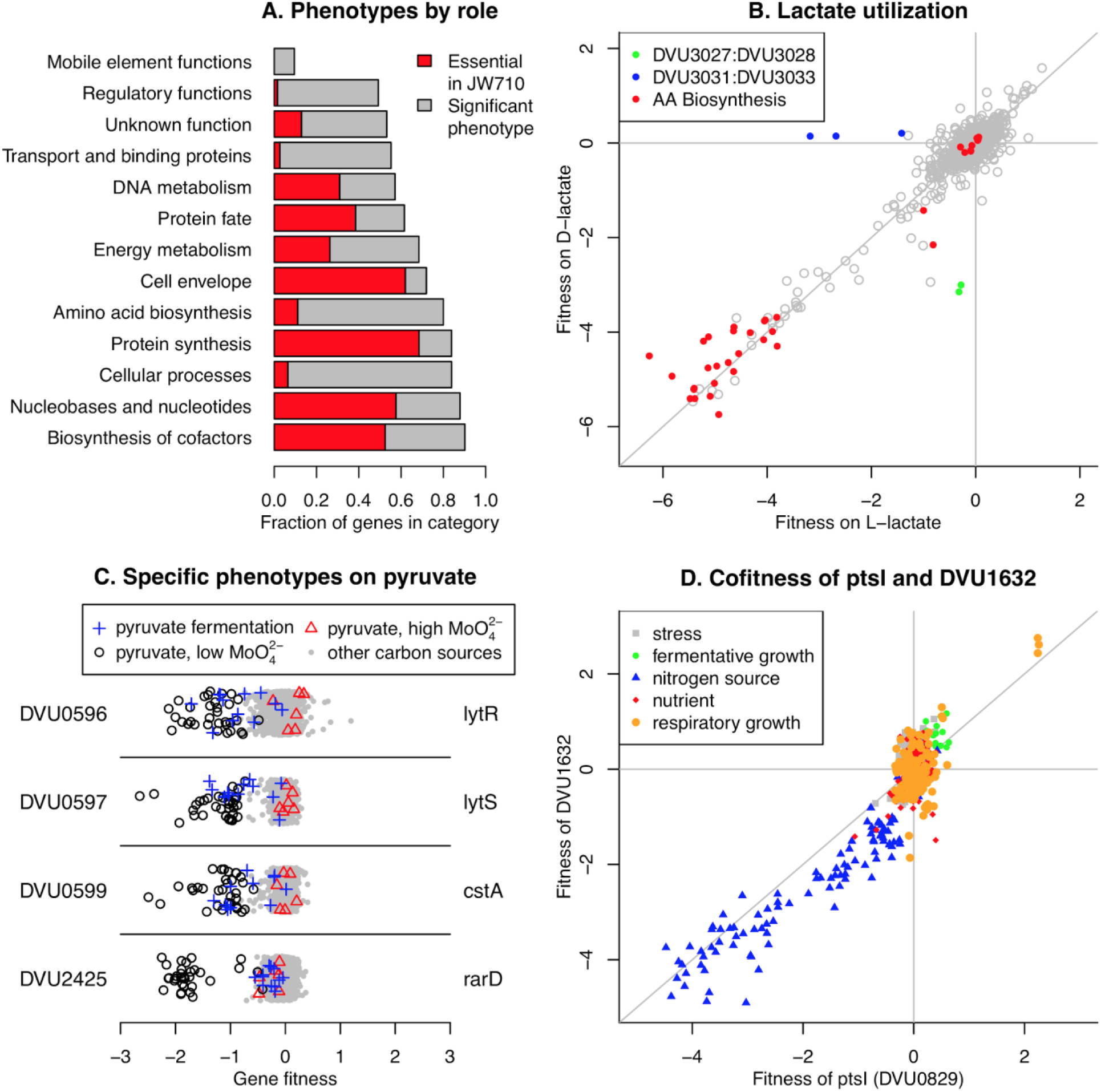
Summary of *D. vulgaris* Hildenborough mutant fitness dataset. (A) Fraction of genes from different functional categories that were essential (in both wild-type and JW710 backgrounds) or had a significant phenotype in at least one experiment. The functional categories are from TIGRFAMs roles (Haft et al., 2013). Only categories with 20 or more genes are shown. (B) Comparison of gene fitness values for growth in minimal media with either L-lactate (*x*-axis) or D-lactate (*y*-axis) as the carbon source. The data is the average of three biological replicates for each condition. “AA biosynthesis genes” are genes with TIGRFAMs role of “Amino acid biosynthesis.” (C) Specific phenotypes for genes important for growth on pyruvate. In these plots, each point represents the fitness of that gene in one of 757 genome-wide assays. Certain experiment classes are highlighted. The *y*-axis is random. (D) Comparison of fitness values for *DVU0829* (*x*-axis) and *DVU1632* (*y*-axis) across all 757 experiments. In panels B and D, lines show *x* = 0, *y* = 0, and *x* = *y*.

Among the 19 genes uniquely essential in the wild-type background (and not in the JW710 background), we did not find a clear biological pattern. We suspect that most of these genes are nearly essential regardless of genetic background, because 18 of the 19 are near our essentiality threshold in the JW710 data (Materials and Methods). Among the 56 genes uniquely essential in JW710, we found that the entire eight gene *de novo* UMP biosynthesis pathway was essential, while all of these genes are clearly dispensable in the wild-type background. These synthetic lethality results are consistent because DvH has two pathways to make UMP, a *de novo* pathway and a salvage pathway (Supplementary Figure 1). Due to the absence of the salvage pathway in JW710 (through the Δ*upp* mutation), the *de novo* pathway becomes essential. Other differences in gene essentiality between JW710 and wild-type DvH in our study, or between gene essentiality in wild-type DvH in our study or in Fels et al. (Fels et al., 2013), could be affected by sequence differences between these strains. Laboratory-acquired mutations in DvH can lead to large phenotypic differences between closely-related strains (De León et al., 2017). To identify genetic variants, we sequenced the JW710 genome and compared these data to our transposon mapping data for our wild-type strain (Materials and Methods). A list of the genetic variants in our wild-type and JW710 strains relative to the reference genome is contained in Supplementary Table 2.

### A gene-phenotype map of DvH

RB-TnSeq simplifies the measurement of mutant phenotypes across multiple experiments through deep sequencing of DNA barcodes that uniquely mark each strain in the library (Price et al., 2018; Wetmore et al., 2015). To facilitate the generation of a large DvH gene-phenotype map, we first constructed a RB-TnSeq mutant library in the JW710 (Δ*upp*) strain. We chose JW710 as the base strain because this background can be used for introducing a second mutation into the library, thus enabling future studies of genetic interactions in DvH. Using Tn-seq, we linked 74,923 unique DNA barcodes to transposon insertions in the main chromosome, and 4,737 unique DNA barcodes to insertions in the DvH native plasmid, pDV1. On both the chromosome and megaplasmid, *Tn*5 insertions were relatively evenly distributed (Supplementary Figure 2). To perform genome-wide mutant fitness assays, we compare the abundance of DNA barcodes after growth selection (referred to as the condition sample) versus before (referred to as the Time0 sample), represented as a log_2_ ratio. To calculate gene fitness scores, we use the weighted average of the individual strain fitness values for mutants in that gene (Wetmore et al., 2015). In the JW710 library, we used 15 independent insertion strains to calculate gene fitness scores for the median gene. Negative gene fitness scores mean that the mutations in this gene made mutants less fit than the average strain in the library, while positive gene fitness scores indicate that the mutations in the gene were beneficial to the mutant in these growth conditions.

Using the JW710 RB-TnSeq library, we generated a large DvH gene-phenotype map by performing 757 genome-wide fitness assays that passed our quality control metrics (Wetmore et al., 2015). In each of these experiments, we assayed the fitness of 2,741 protein-coding genes, for a total of 2.07 million gene-phenotype measurements. (241 non-essential proteins do not have fitness values, most often because mutants in these genes are at low abundance in the Time0 samples.) To systematically investigate the physiology of DvH, we assayed a diverse range of conditions including respiratory growth, fermentative growth, growth in the presence of different nutrients, and growth in the presence of different stressors. The complete list of experiments, with associated metadata, is available in Supplementary Table 3. These 757 experiments include 244 unique experimental conditions (the remainder are biological replicates). The DvH fitness dataset can be explored interactively at the Fitness Browser (fit.genomics.lbl.gov), which bundles multiple computational tools to aid in the elucidation of novel gene functions. To illustrate the data, we highlight a comparison of two conditions, growth in defined media with either L-lactate or D-lactate as the sole carbon source and electron donor. As expected, many genes involved in the biosynthesis in amino acids had large fitness detects in both conditions (Figure 1B), while only a few genes had phenotypes unique to each substrate. In support of previous observations (Vita et al., 2015), our data demonstrates that DVU3032:DVU3033 encodes L-lactate dehydrogenase while DVU3027:DVU3028 encodes D-lactate dehydrogenase, as these genes have large growth defects on each substrate (Figure 1B). In addition, we found that DVU3031 is also important for growth on L-lactate. DVU3031 encodes a conserved but experimentally uncharacterized protein with AAA and DRTGG domains, and our data provides the first experimental evidence of the importance of this gene for growth on L-lactate, although its precise function remains to be determined.

Across the entire fitness dataset, we identified a significant phenotype (|fitness| > 0.5 and |*t*| > 4, where *t* is a measure of the significance of the measurement (Wetmore et al., 2015)) for 1,137 genes in at least one experiment (at an estimated false discovery rate of 3%; Materials and Methods). While non-essential genes from all functional categories (main roles from TIGRFAMs (Haft et al., 2013)) had significant phenotypes, those involved in general cellular processes, amino acid biosynthesis, and transport were more likely to have a phenotype in one of our experiments, while those involved in mobile and extrachromosomal element functions, regulation, and the cell envelope were less likely to have a phenotype (Figure 1A). Among non-essential proteins with vague or hypothetical annotations, we identified a conditional phenotype for 34%. Our dataset provides a starting point for uncovering the roles for these genes.

In our prior work, we have used two primary strategies to infer gene functions from mutant phenotypes: specific phenotypes and cofitness (Price et al., 2018). A gene has a specific phenotype if it has a phenotype in only one or a handful of conditions (Materials and Methods), in contrast to genes with more pleiotropic effects. Intuitively, specific phenotypes are informative for gene function because hypotheses can be readily derived from the one or few conditions where a phenotype is observed. For example, we found that *DVU0599* (*cstA*) had a specific phenotype under conditions where pyruvate was used as the sole carbon source (Figure 1C), suggesting that this gene is involved in pyruvate utilization. DVU0599 is distantly related to *Escherichia coli* YjiY (32% amino acid identity), which was recently demonstrated to be a pyruvate transporter (Kristoficova et al., 2018). The pyruvate-specific phenotype of *DVU0599* strongly suggests that it also encodes a pyruvate transporter. We also identified pyruvate-specific phenotypes for the nearby two component signaling system encoded by *DVU0596:DVU0597* (Figure 1C), which is consistent with the known regulation of *DVU0599* by this system (Rajeev et al., 2011). Lastly, we found a similar pyruvate-specific phenotype for *DVU2425* (*rarD*), which encodes an uncharacterized protein conserved in diverse bacteria (Figure 1C). RarD-family proteins are predicted to be transporters, and thus it is possible that DVU2425 also transports pyruvate. However, in other bacteria with available fitness data (Price et al., 2018), RarD proteins are important for transporting various amino acids. Across the entire dataset, we identified specific phenotypes for 540 different genes. These specific phenotypes are spread across a wide range of different experimental conditions, suggesting that they are useful for understanding different aspects of DvH biology.

Genes that have high cofitness (*r*, correlated patterns of phenotypes across all experiments) are likely to share a cellular function (Deutschbauer et al., 2011; Price et al., 2018). For example, the phosphotransferase system (PTS) proteins PtsI (DVU0829) and DVU1632 (a putative EII-A enzyme) are highly cofit (*r* = 0.89), with both genes sharing fitness defects in a number of experiments with alternative nitrogen sources (Figure 1D). In the entire DvH dataset, we identified 2,104 gene pairs that have high cofitness (*r* >= 0.8), with 330 different genes having at least one cofitness relationship. In the subsequent sections, we combine the specific phenotypes and cofitness relationships with comparative genomics to derive new insights into the functions of poorly understood DvH genes.

### *DVU0867* encodes an atypical L-aspartate decarboxylase

To identify genes that are required for vitamin biosynthesis, we tested growth in minimal medium with vitamins omitted. (Our defined medium usually contained Thauer’s vitamins.) To stimulate growth and hence vitamin requirements, we added vitamin-free casamino acids (a mixture of amino acids) to the media for many of these experiments. We will describe novel findings in the biosynthesis of pantothenic acid (vitamin B_5_), biotin (vitamin B_7_), and pyridoxal phosphate (vitamin B_6_).

First, as shown in Figure 2A, we identified three genes that were specifically important when pantothenic acid was not available. Two of these genes (*DVU2446* and *DVU2448*) were already annotated as being involved in pantothenic acid biosynthesis (*panB* and *panC*, respectively). The remaining gene, *DVU0867*, was annotated as aromatic amino acid decarboxylase. DVU0867 is distantly related (27% amino acid identity) to the aspartate decarboxylase (PanP) of *Vibrio fischeri (Pan et al*., *2017)*. Aspartate decarboxylase is expected to be required for synthesis of pantothenic acid (Figure 2B). *E. coli* and many other bacteria encode *panD* for this step, but neither *panD* nor its maturation cofactor *panZ* are present in DvH genome. So, we hypothesized that *DVU0867* encodes an aspartate decarboxylase.

To test if DVU0867 could decarboxylate L-aspartate, we attempted to complement mutants of *E. coli* that require pantothenic acid for growth. The expression of *DVU0867* restored growth of the *panD* mutant in the absence of pantothenic acid (Figure 2C), but did not restore growth of *panC* or *panB* mutants (data not shown). This confirmed that the protein encoded by *DVU0867* performs the same function as PanD, L-aspartate decarboxylase.

### Biotin synthesis with a specialized acyl carrier protein and without methyl esters

Biotin, commonly known as vitamin B_7_, is an essential enzyme cofactor required by all three domains of life. However, it remains unclear how DvH synthesizes biotin, as it does not contain clear homologs for the entirety of known biosynthetic pathways. Because biotin is only required in trace amounts, we added the protein avidin (0.1 U/mL), which has a high affinity for biotin and sequesters it, to our no-biotin experiments. We identified ten genes that were specifically important for growth in the absence of biotin (Figure 3A). Nine of these ten genes were clustered together (*DVU2558:DVU2565*). This cluster includes *bioF, bioA, bioD*, and *bioB*, which together convert pimeloyl-[ACP] to biotin. (ACP is short for acyl carrier protein.) These genes bracket a cluster of genes encoding homologs to known fatty acid biosynthesis factors: DVU2560 and DORF42491 have FabZ-like domains, DVU2561 is a putative 3-oxoacyl-ACP reductase, DVU2562 is homologous to acyl-carrier protein, and DVU2563 contains a beta-keto-acyl carrier protein synthase (KAS) domain and is annotated as a FabF protein. These genes have previously been suggested to play a role as an alternate pathway for the synthesis of pimeloyl-ACP from malonyl-CoA, which is the first stage of *de novo* biotin synthesis (Lin and Cronan, 2011; Rodionov et al., 2004). The phenotype observed in our fitness assays are the first experimental evidence supporting their role in synthesizing biotin. Another gene in the primary biotin synthesis cluster, *birA* (*DVU2557*) encoding a transcriptional repressor and biotin-protein ligase, did not have a strong phenotype in our no-biotin experiments. This is possibly due to the presence of another putative biotin-protein ligase (DVU1835) in the genome.

**Figure 2.**
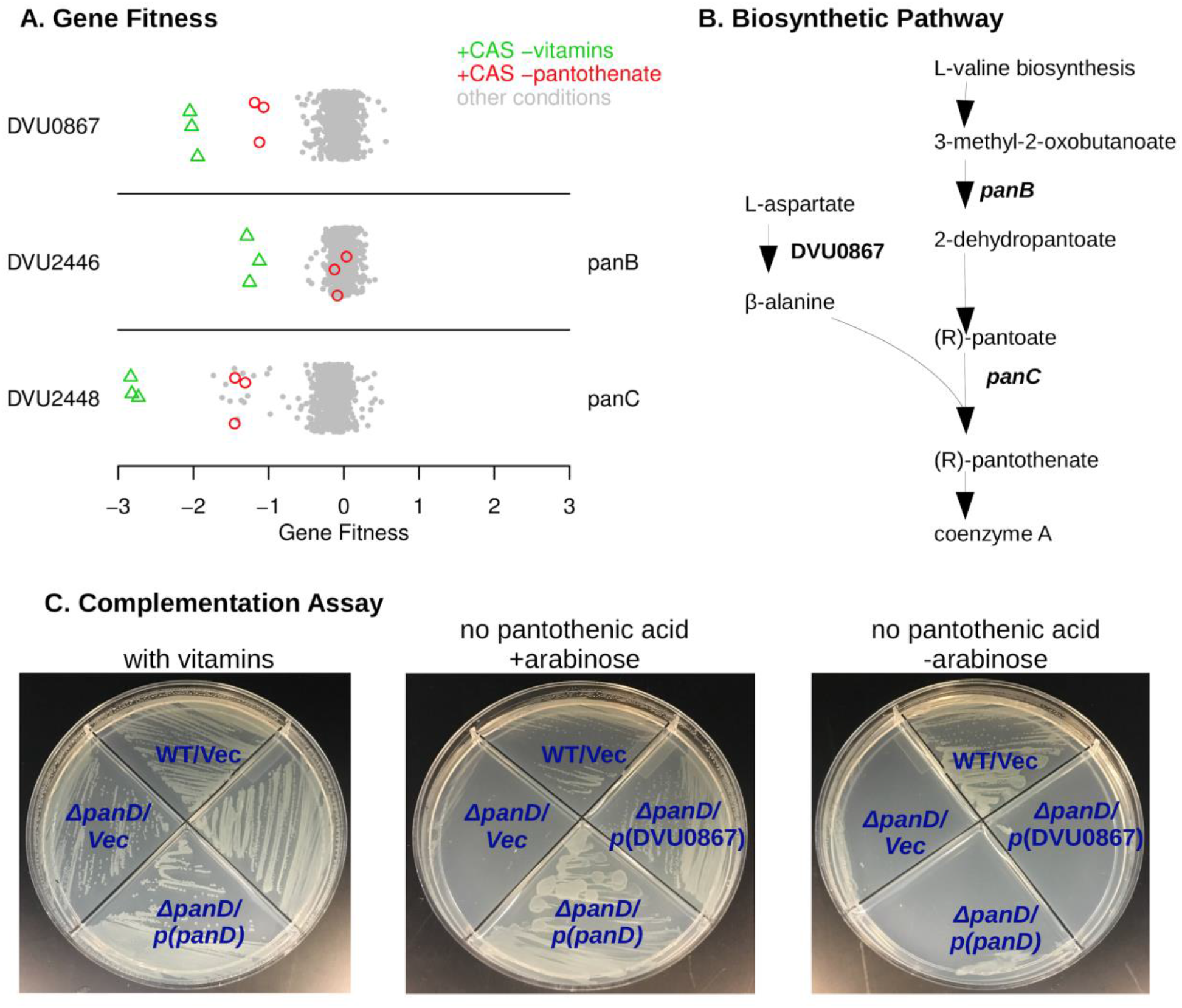
DVU0867 has L-aspartate 1-decarboxylase activity. (A) Fitness data for each gene across all 757 experiments, with pantothenate-related conditions highlighted. The *y*-axis is random and CAS is short for casamino acids. (B) The biosynthetic pathway for pantothenate, with key enzymes highlighted. (C) Complementation assays using an *E. coli panD* gene knockout strain that is defective in pantothenate biosynthesis and lacks aspartate 1-decarboxylase activity. Indicated genes were cloned upstream of an arabinose inducible promoter. Vec is an empty vector control.

**Figure 3.**
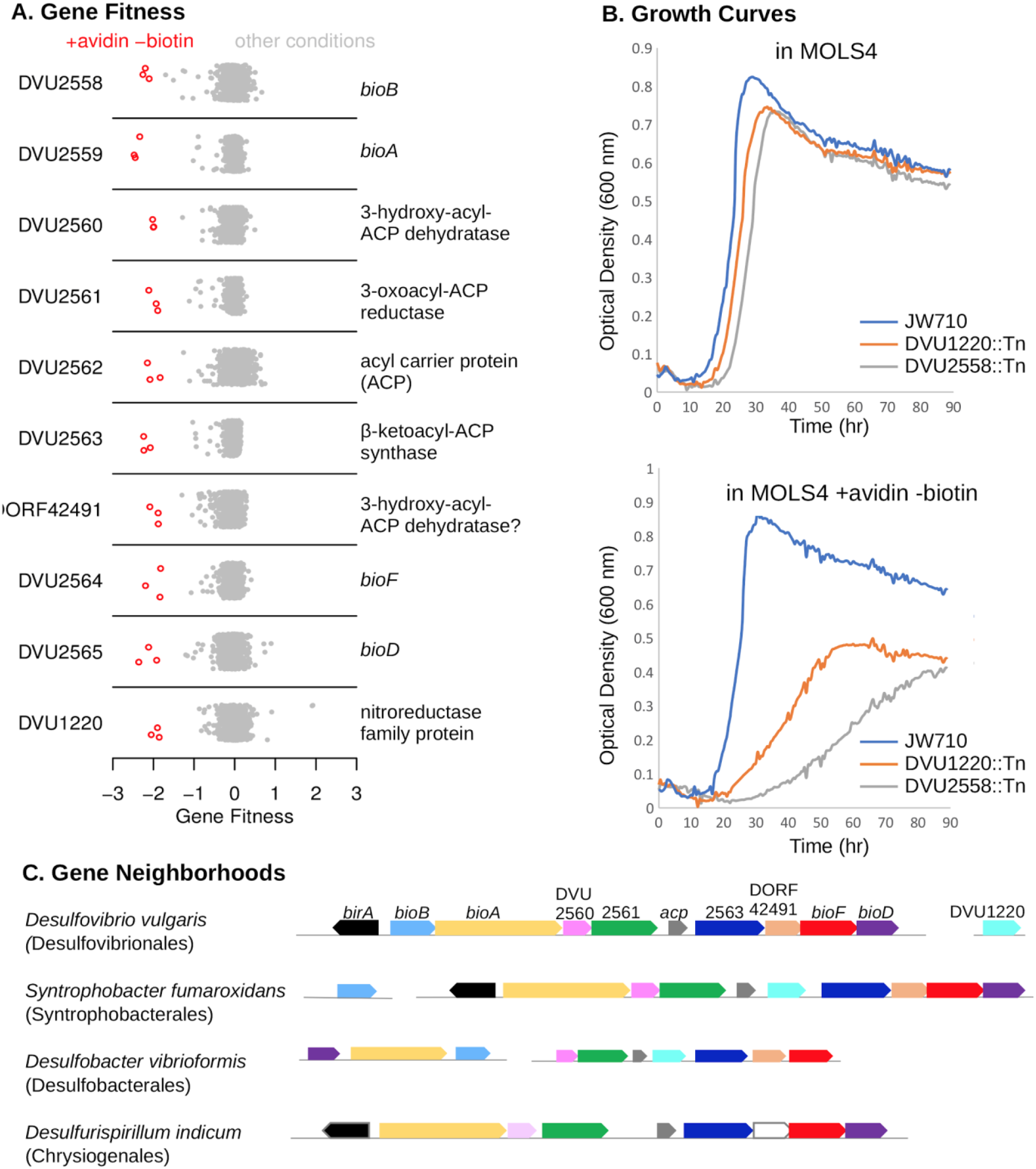
An atypical biotin synthesis pathway in *D. vulgaris* Hildenborough. (A) Fitness data for each gene across all 757 experiments, with experiments in the absence of biotin and supplemented with avidin highlighted. The *y*-axis is random. (B) Growth assays of the DvH JW710 control strain, and strains with mutations in either *DVU1220* or *DVU2558* (*bioB*). Measurements were made in a Bioscreen C growth analysis system and each curve is the average of four replicates. (C) Gene neighborhood and conservation in other microorganisms with components of the DvH biotin synthesis pathway. Genes are colored by their homology to the corresponding DvH genes, and *birA* is only shown if it is adjacent to other biotin synthesis genes.

The single gene outside of this gene cluster whose mutant displayed a specific fitness defect in the absence of biotin was *DVU1220*, which is annotated as a nitroreductase. DVU1220 is predicted to contain both flavin mononucleotide (FMN) and [4Fe4S] iron sulfur cluster cofactors. We monitored the growth of the parental strain, JW710, alongside *DVU1220* and *DVU2558* (*bioB)* mutants in media with and without biotin depletion (Figure 3B). Both *DVU1220* and *DVU2558* mutants displayed significant growth defects under biotin depletion conditions, which confirms a role for *DVU1220* in biotin synthesis.

The existence of an extended biotin gene cluster was previously reported to be limited to the genus *Desulfovibrio* (Rodionov et al., 2004); (Lin and Cronan, 2011). More genome sequences are now available and we found similar gene clusters in two other orders of Deltaproteobacteria and in *Desulfurispirillum indicum* from the phylum Chrysiogenetes (Figure 3C, Supplementary Figure 3). Furthermore, in the genomes of *Syntrophobacter fumaroxidans* MPOB and *Desulfobacter vibrioformis* DSM 8776 for instance, these gene clusters include a nitroreductase-like gene (Figure 3C, Supplementary Figure 3). The nitroreductases in these biotin clusters are similar to each other (over 40% pairwise identity) but are distantly related to DVU1220; nevertheless, their clustering with the other genes of the pathway is consistent with the involvement of a nitroreductase-like protein in biotin synthesis in these organisms.

Although our data show that the entire biotin synthesis cluster as well as *DVU1220* are required for biotin synthesis in DvH, further study will be needed to define the biochemical pathway for the first stage of biotin synthesis, up to pimeloyl-ACP. It appears that *DVU2562* encodes an alternate ACP that is specialized for this pathway. (The other ACP in the genome, *DVU1205*, is essential, presumably because it is required for fatty acid biosynthesis.) In contrast to *E. coli*, where the enzymes for fatty acid biosynthesis convert malonyl-ACP methyl ester to pimeloyl-ACP methyl ester, it appears that DvH uses specialized enzymes to elongate malonyl-ACP to pimeloyl-ACP, without methyl ester intermediates. Indeed, neither the gene for forming the methyl ester (*bioC*) nor the gene for removing it (*bioH*) are found in DvH. The elongation of malonyl-ACP to pimeloyl-ACP would require a β-keto-ACP synthase, a 3-oxo-ACP reductase, a β-hydroxyacyl-ACP dehydratase, and a enoyl-ACP reductase. The cluster contains candidates for all of these activities except for enoyl-ACP reductase, which might be provided by the nitroreductase-like protein or by a promiscuous enzyme from fatty acid biosynthesis (such as DVU2064, which is essential).

### The putative dehydrogenase DVU0826:DVU0827 is required for vitamin B_6_ synthesis

By growing the mutant pool in defined medium that lacks pyridoxal phosphate (vitamin B_6_), we identified a putative two-subunit dehydrogenase (DVU0826 and DVU0827) that is required for pyridoxal phosphate biosynthesis (Figure 4A). The genes for both subunits have nearly identical fitness patterns (high cofitness) as *pdxA* (*DVU2241*), which encodes 4-hydroxythreonine-4-phosphate dehydrogenase. Besides *pdxA* and the dehydrogenase, the DvH genome also encodes pyridoxine 5’-phosphate synthase (*pdxJ, DVU1908*), which implies that DvH synthesizes pyridoxal phosphate via deoxyxylulose 5’-phosphate, as in *E. coli* (Figure 4B) (Mittenhuber 2001; Fitzpatrick et al., 2007). The orthologs of DVU0826:DVU0827 in *D. vulgaris* Miyazaki F (DvMF_2874:DvMF_2875) display their highest co-fitness values (*r* = 0.78 and 0.87, respectively) with PdxJ (DvMF_0281) (data from Price et al 2018). This evidence supports a role for the dehydrogenase in pyridoxal phosphate biosynthesis.

**Figure 4.**
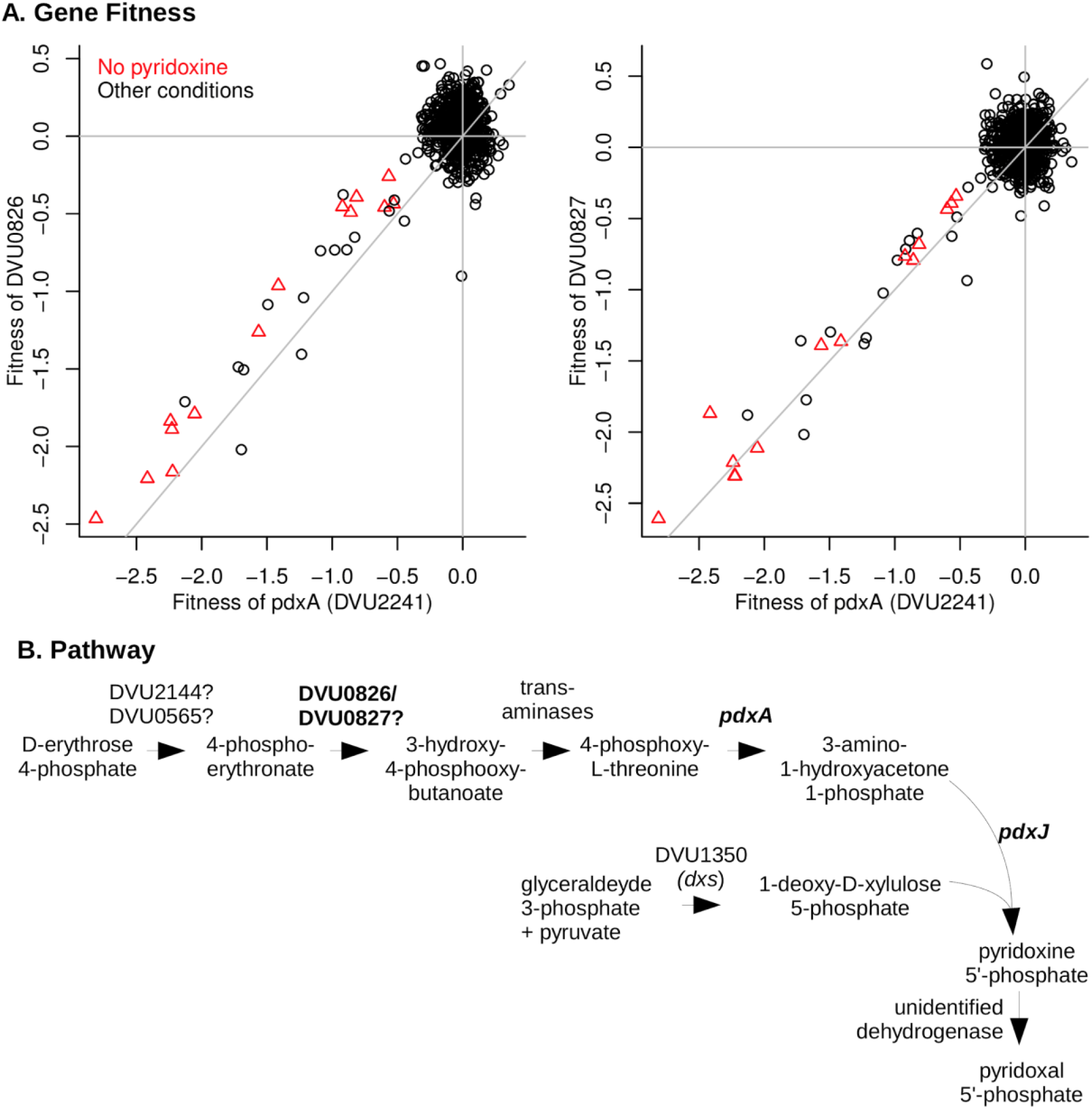
*DVU0826* and *DVU0827* are required for vitamin B6 synthesis. (A) Comparison of gene fitness values across 757 experiments between *pdxA* (*DVU2241*) and either *DVU0826* (left) or *DVU0827* (right). Experiments performed in the absence of pyridoxine are highlighted. *pdxA* has high cofitness with both *DVU0826* (*r* = 0.81) and *DVU0827* (*r* = 0.84). (B) The proposed pathway of pyridoxal phosphate biosynthesis in DvH.

As shown in Figure 4B, the DvH genome seems to be missing genes for two dehydrogenase enzymes in pyridoxal phosphate biosynthesis: 4-phosphoerythronate dehydrogenase (PdxB in *E. coli)* and pyridoxine 5’-phosphate oxidase (PdxH in *E. coli). DVU0921* encodes a pyridoxamine 5’-phosphate oxidase domain (PF12900), but this putative protein is very distantly related to PdxH, and insertions in *DVU0921* exhibited little phenotype in any of our assays (all |fitness| < 1), so we do not think it encodes the missing PdxH. *PdxH* is essential for the growth of most bacteria in media that contain yeast extract (data of Price et al 2018) because PdxH is required to convert pyridoxine to pyridoxal phosphate. Since *DVU0826:DVU0827* are not essential, we suspect that they encode a novel 4-phosphoerythronate dehydrogenase.

### Utilization of nitrogen sources

DvH is capable of fixing nitrogen gas (Heidelberg et al., 2004; Riederer-Henderson and Wilson, 1970) but its capacity to use other nitrogen sources in the presence of pDV1-encoding nitrogenase has not been reported. As far as we know, DvH does not use amino acids as the sole source of carbon for growth. We assayed gene fitness for DvH in defined lactate-sulfate media with 28 different nitrogen sources, including with N_2_ only. As expected, mutations in the *nifD* and *nifK* genes, encoding the alpha and beta subunits of the nitrogenase complex respectively, were highly detrimental to growth when N_2_ was the sole nitrogen source available, but had little effect on growth in ammonium (Figure 5). All of the genes in the nitrogen fixation cluster (*DVUA0007:DVUA0016*) were important for growth with no added nitrogen, except for *DVUA0010*, which had no fitness data. (After averaging across six replicate experiments with N_2_ as the nitrogen source, each other gene in the cluster had fitness < −2.)

**Figure 5.**
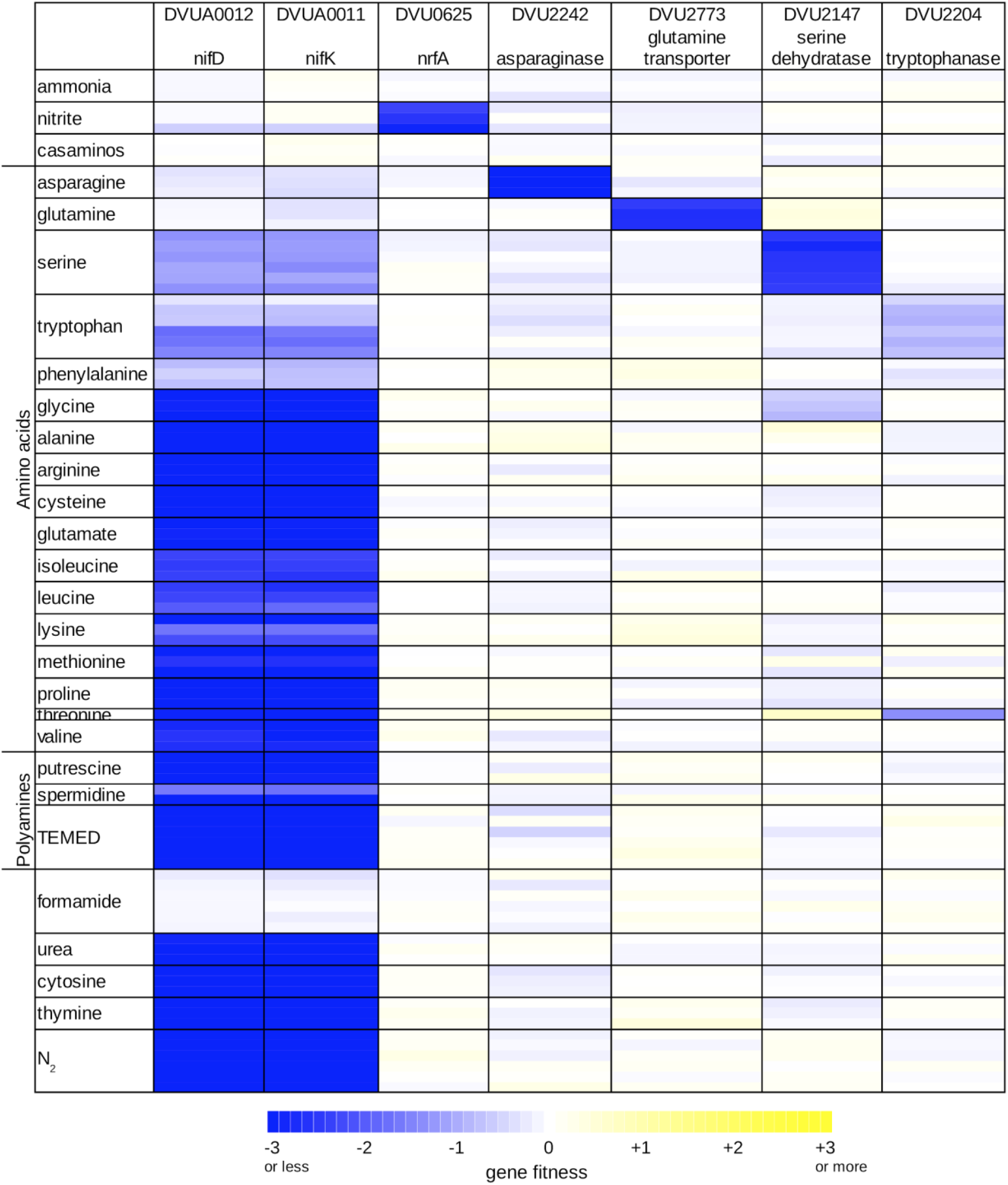
Overview of nitrogen utilization in *D. vulgaris* Hildenborough. Heatmap of gene fitness data for select genes in experiments where the nitrogen source was varied. For each condition, we show the data from each replicate experiment separately. TEMED is tetramethylethylenediamine; casaminos is casamino acids.

We used the fitness of the nitrogenase genes *nifDK* to identify additional conditions under which nitrogen fixation contributed to growth. We considered the nitrogen sources we tested as well-utilized if *nifDK* did not contribute to fitness, and as weakly utilized if *nifDK* had a milder phenotype. Based on the fitness data of *nifDK*, we found that among the amino acids glutamine and asparagine were well-utilized, and that serine, tryptophan, and phenylalanine were weakly utilized (Figure 5). In addition, we found that growth in minimal medium with no ammonium was stimulated by the addition of glutamine, asparagine, serine, or phenylalanine (data not shown), which further confirms that these amino acids are utilized as nitrogen sources by DvH. No growth after five days was observed when histidine, aspartate, or tyrosine were provided as alternatives to ammonium. Growth occurred in the presence of the remaining 11 amino acids, but appears to depend entirely on nitrogen fixation (Figure 5), which suggests that they are not utilized by DvH as nitrogen sources. We did find that the addition of valine, isoleucine, leucine, or glutamate slightly reduced the initial lag phase when DvH was grown in minimal medium with no ammonium (Supplementary Figure 4). Nevertheless, because nitrogen fixation genes were important in these conditions, and because we did not identify potential catabolic genes or transporters that were important for utilizing these four amino acids, we believe that they are not utilized. The simulation of growth could be due to effects on gene regulation or due the uptake of small amounts of amino acids.

The strong fitness defect of mutants in the asparaginase encoded by *DVU2242*, when asparagine was used as sole nitrogen source (Figure 5), shows that this enzyme can efficiently provide ammonium. We did not identify any enzymes that were specifically important during growth on glutamine, so the origin of the glutaminase activity remains unclear. We did identify a putative transporter of the NbcE family (TC 2.A.115, (Saier et al., 2016)) that was specifically important for growth on glutamine (DVU2773), so we propose that *DVU2773* encodes the glutamine transporter. The putative serine dehydratase (*DVU2147*) and tryptophanase (*DVU2204*) were important for growth on serine and tryptophan, respectively (Figure 5), which confirms their participation in providing ammonium from these amino acids. We are not sure why both nitrogenase and a deaminating enzyme were important for growth with serine or tryptophan. It is possible that uptake is slow, that the deaminating enzymes are weakly expressed, or that nitrogen fixation and deamination are important during different phases of growth. We did not identify any genes that were specifically important for utilizing phenylalanine.

In addition to amino acids, we tested the utilization of polyamine, nucleobases, urea, nitrite, and formamide. In the presence of nitrite or formamide, nitrogen fixation was not required (Figure 5), which shows that DvH can utilize these nitrogen sources as well. As expected, nitrite utilization required the nitrite reductase NrfA (DVU0625; Figure 5). Although the DvH genome contains a gene annotated as a formamidase (*DVU1164*), we did not identify any phenotypes for this gene (all |fitness| < 0.5). Overall, we found that DvH can utilize nine nitrogen sources (ammonia, N_2_, five of the amino acids, nitrite, and formamide), and we identified genes involved in the utilization of most of these nitrogen sources.

### Chlorate toxicity is mediated via the aldehyde oxidoreductase (Aor)

The use of chlorate has been proposed as an additive to control the growth of SRB and concomitant sulfide production, which causes oil souring and is a major industrial problem (Engelbrektson et al., 2014; Gregoire et al., 2014). Previous work indicated that (per)chlorate can serve as specific and potent inhibitors of sulfate respiration (Carlson et al., 2015). It has been shown that both perchlorate and chlorate act, in part, as direct competitive inhibitors of sulfate adenylyltransferase, the first step in the pathway (Carlson et al., 2015; Mehta-Kolte et al., 2019; Stoeva and Coates, 2019). Alternatively, reduction of (per)chlorate and their conversion into reactive chlorine species (RCS) (chlorite and hypochlorite) has been attributed to the adventitious reactivity of metal-binding pterin dependent enzymes such as nitrate reductase, which has a molybdenum cofactor.

Although the DvH genome does not contain a predicted (per)chlorate reductase or nitrate reductase, the growth of DvH JW710 was significantly reduced when cultured in the presence of 10 mM chlorate and was almost completely inhibited upon addition of 20 mM chlorate (Figure 6A). Fitness profiling the chemogenomic response of JW710 to 6.25 mM chlorate identified 15 genes that were detrimental to fitness (gave a positive fitness value when disrupted) in this condition but had little effect on fitness in the absence of chlorate (Figure 6A). Seven of these genes are involved in the biosynthesis of molybdenum cofactor or tungsten cofactor. (These are *moaA* (*DVU0580*), *moaC* (*DVU0289*), *moaE* (*DVU2212*), *moeA* (*DVU2990*), *moeA-2* (*DVU0951*), *moeB* (*DVU0643*), and *mogA* (*DVU0971*)). Another four genes are involved in the uptake of tungstate (*tupABC* or *DVU0747:DVU0745*) or its regulation (*tupR* or *DVU3193*; (Rajeev et al., 2018)). The gene with the largest effect on fitness was *aor* (*DVU1179*), encoding a putative tungsten-dependent aldehyde:ferredoxin oxidoreductase.

**Figure 6.**
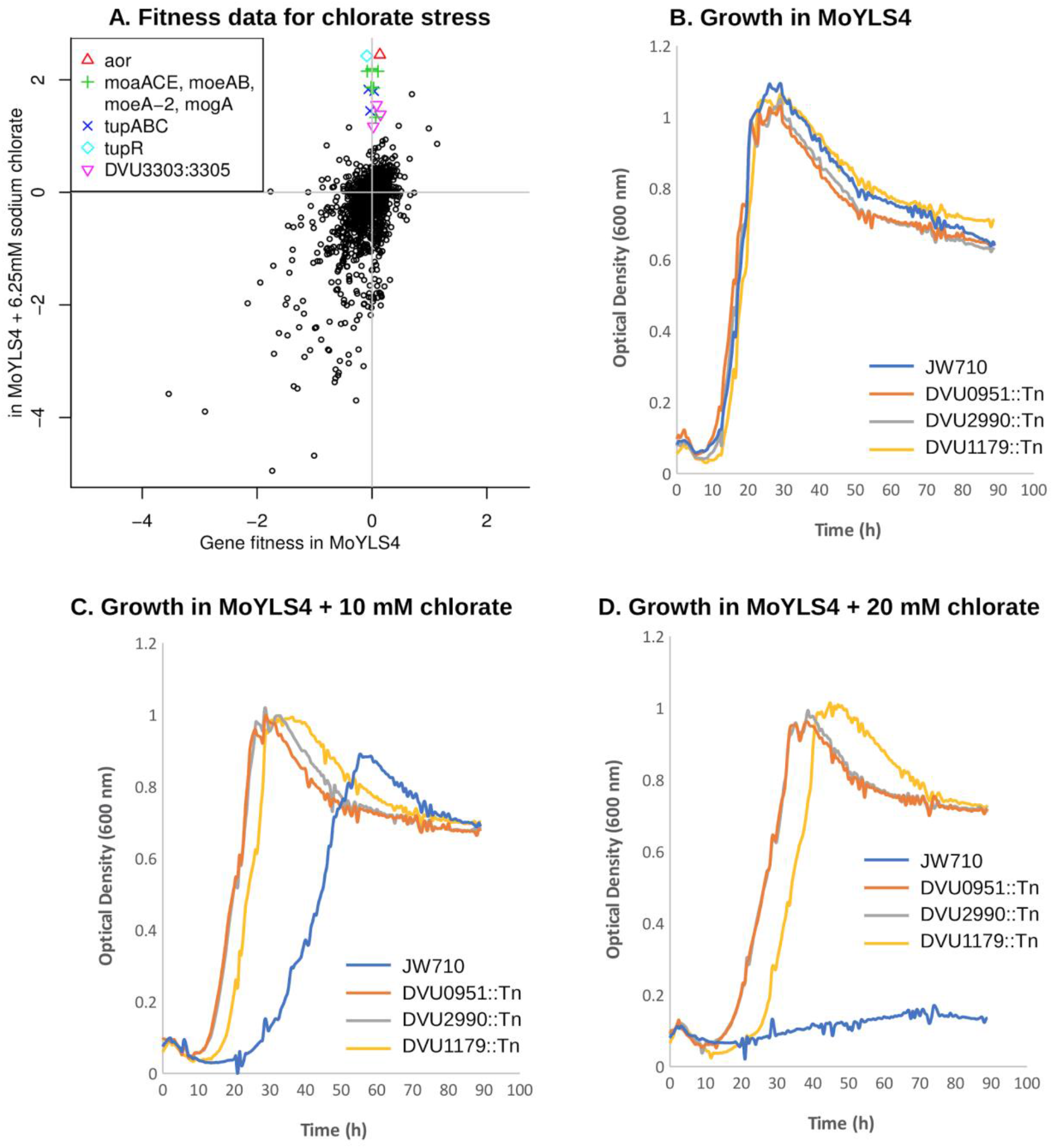
Loss of aldehyde oxidoreductase activity results in chlorate resistance. (A) Comparison of gene fitness values for growth in rich lactate-sulfate media or in media supplemented with 6.25 mM sodium chlorate. Each value is the average from three replicate experiments. (B-D) Growth of DvH JW710 and mutant strains of DVU0951 (*moeA*), DVU2990 *(moeA-2)* and DVU1179 (*aor*) in rich media with increasing concentrations of sodium chlorate. Each curve is the average of four replicates.

It appears that biosynthesis of the tungsten cofactor is detrimental to the parental strain in the presence of chlorate because it allows for the activity of Aor. Furthermore, this effect does not occur with other oxyanions of chlorine: mutants in *aor* and the other tungsten-related genes were about as sensitive as other mutants in the presence of perchlorate or chlorite. (Fitness values for all twelve of those genes when challenged with 6.25-12.5 mM perchlorate or 0.1-0.25 mM chlorite were between −1 and +1 across all three replicates of each condition.) To quantify the advantage of disrupting *aor* or tungsten cofactor biosynthesis genes during growth on chlorate, we compared the growth of mutant strains for *aor, moeA*, and *moeA-2* to the library parental strain JW710 in presence of up to 20 mM chlorate. All strains grew similarly in the absence of chlorate (Figure 6B); JW710 was inhibited by 10 mM chlorate (Figure 6C); and the interruption of *aor* or of the tungsten cofactor biosynthesis genes conferred resistance to 20 mM chlorate (Figure 6D).

Based on these data, we propose that Aor catalyzes the reduction of chlorate to chlorite, which is far more toxic. Thus, disruption of *aor* itself, or of various genes involved in the acquisition of tungstate or the biosynthesis of tungsten cofactor, will confer resistance to chlorate. To test if this mechanism of chlorate toxicity applies to other sulfate-reducing bacteria, we examined previously-published fitness data for *D. vulgaris* Miyazaki F and *D. alaskensis* G20 growing in lactate-sulfate medium in the presence of chlorate. In *D. vulgaris* Miyazaki F, the three most detrimental genes during growth in 6.25 mM chlorate were all involved in molybdenum or tungsten cofactor biosynthesis (fitness > +6.0, data of (Price et al., 2018)). The ortholog of *aor* (*DvMF_1956*) showed strongly positive fitness (fitness = +5.6) and two other molybdopterin-containing enzymes predicted to be anaerobic dehydrogenases (*DvMF_1484* and *DvMF_0448*) had milder positive fitness values (fitness = +2 and +1.5 respectively). This suggests that several enzymes contribute to the reduction of chlorate in *D. vulgaris* Miyazaki F. In *D. alaskensis* G20, the six genes providing functions that were most detrimental to growth in the presence of chlorate are all involved in molybdenum or tungsten cofactor biosynthesis (data of (Carlson et al., 2015)). Thus, in other sulfate-reducing bacteria, the activity of molybdopterin-dependent enzymes is involved in chlorate toxicity.

Our data also allow the inference that both *moeA* and *moeA*-2 are required for the formation of the tungsten cofactor of *aor*. Furthermore, the two genes have high cofitness across all of our experiments (*r* = 0.85). Their orthologs in *D. vulgaris* Miyazaki F (*DvMF_1797* and *DvMF_1358*) also have high cofitness (*r* = 0.90). The two *moeA*-like proteins of DvH are distantly related (33% identity) and DVU0951 (annotated as MoeA) has an additional C-terminal domain (PF12727) from the periplasmic binding protein superfamily. It is not understood why many bacterial genomes contain two *moeA*-like genes, unlike *E. coli*, which has a single *moeA*. Some researchers have speculated that the two copies might be specialized for the insertion of molybdate or tungstate (i.e., (Smart et al., 2009)), but this is not the case in *Desulfovibrio*, nor are the two *moeA* genes functionally redundant since the inactivation of each resulted in resistance to chlorate.

The remaining three genes that are detrimental during growth of JW710 in 6.25 mM chlorate form a putative three-component signaling system that includes a Lon-type protease, a histidine kinase, and a DNA-binding response regulator (DVU3303:DVU3305). The three genes have very similar fitness patterns (all pairwise cofitness > 0.9), which confirms that they function together. One of the regulatory targets of this system is the putative anion transporter encoded by *DVU3299* (Rajeev et al., 2011). *DVU3299* is important for growth in chlorate stress (average fitness = −2.3 across three replicates), so the phenotype of the signaling system may be due to its effect on the expression of *DVU3299*.

### The response to molybdate toxicity

Molybdate (MoO_4_^2-^) is highly toxic to SRB due to its capacity to act as a futile substrate for the adenosine phosphosulfate (APS) reductase, a key enzyme in the sulfate reduction pathway (Peck, 1962). We examined the response of the JW710 mutant pool when challenged with 100 uM Na_2_MoO_4_ under lactate-sulfate, pyruvate-sulfate and pyruvate-sulfite growth conditions using BarSeq. A total of 40 genes were found to respond specifically to the presence of molybdate. Here we will highlight two gene clusters that were not previously linked to molybdate and which exhibited among the strongest responses: *DVU0436:DVU0438* and *DVU0539:DVU0545* (Figure 7A).

**Figure 7.**
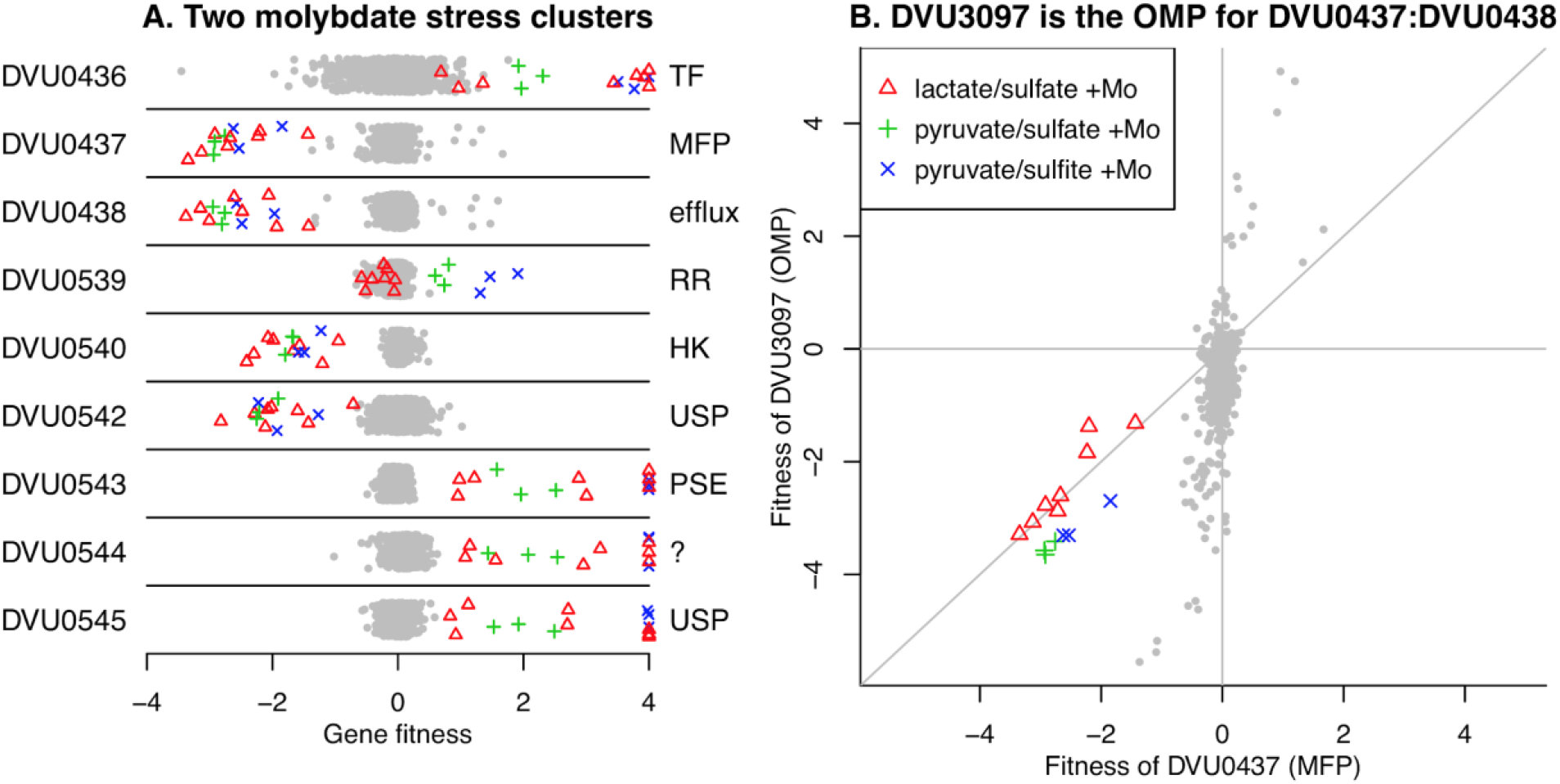
Selected genes with specific phenotypes during molybdate stress. (A) Two clusters of genes involved in molybdate stress. Each point represents the fitness of that gene in a genome-wide assay (*x*-axis). Values above +4 are shown at +4. The *y*-axis is random. (B) Comparison of fitness patterns for the membrane fusion protein DVU0437 and the outer membrane protein DVU3097. In both panels, experiments with added molybdate (0.1 mM) are highlighted with the same color coding (see panel B legend). 4 of the 8 experiments with lactate/sulfate media and added molybdate also had tungstate added (at 0.5 or 2.0 mM).

First, mutations in *DVU0436*, encoding a TetR-type transcriptional regulator, displayed a fitness advantage for the mutants in the presence of molybdate (Figure 7A). In contrast, strains lacking *DVU0437* or *DVU0438*, which are annotated as the membrane fusion protein (MFP) and the integral membrane subunits of a resistance-nodulation-division (RND) type efflux pump, had significant fitness defects in the presence of molybdate (Figure 7A). These data suggest that DVU0436 represses transcription of the RND-type efflux pump encoded by *DVU0437:DVU0438*. Indeed, the RegPrecise database predicts that DVU0436 regulates the operon *DVU0436:DVU0438* via a site just upstream of *DVU0436* (Novichkov et al., 2013).

RND efflux pumps have a third outer membrane component, but no candidates for the missing component were found near *DVU0437:DVU0438*. Instead, based on cofitness, we identified DVU3097 as the probable outer membrane component. *DVU3097* shares the molybdate-related phenotypes of *DVU0437:DVU0438* but also has other phenotypes (Figure 7B). The other phenotypes suggest that *DVU3097* encodes the outer membrane component of other efflux systems as well. The orthologous efflux system in *D. vulgaris* Miyazaki F (*DvMF_1516:DvMF_1515*/*DvMF_2365*) is specifically important for growth in the presence of the antibiotic carbenicillin (data of (Price et al., 2018)). In both strains of *Desulfovibrio*, the efflux system is probably involved in maintaining cell wall integrity rather than in the efflux of molybdate or a related compound.

The second molybdate-responsive cluster, *DVU0539:DVU0545*, is comprised of two divergently transcribed operons, *DVU0540:DVU0539* and *DVU0542:DVU0545*. (There is no *DVU0541* gene.) *DVU0540:DVU0539* encode a sensor histidine kinase and a DNA-binding response regulator, respectively. DVU0539 is thought to regulate both operons, and when cloned into *E. coli*, DVU0539 can activate transcription from *DVU0542*’s promoter (Rajeev et al., 2011). *DVU0542:DVU0545* encode two proteins with homology to universal stress proteins (DVU0542, DVU0545), a transporter from the putative sulfate exporter (PSE) family (DVU0543; TC 2.A.98; (Saier et al., 2016), and a 133-amino acid protein with a transmembrane helix (DVU0544). The histidine kinase (DVU0540) and one of the universal stress proteins (DVU0542) were important for growth in the presence of molybdate, while the other genes in the cluster were detrimental during growth in the presence of molybdate, resulting in a positive fitness value when disrupted (Figure 7A). The only exception was the response regulator (DVU0539) had no phenotype during molybdate stress in lactate-sulfate medium (Figure 7A). Based on these data, it appears that the histidine kinase DVU0540 opposes the activity of the response regulator DVU0539. Nevertheless, the roles of *DVU0542:DVU0545* in responding to molybdate remain unclear and will require further investigation.

## CONCLUSION

Despite the importance of SRB to many environmental and industrial processes, we still have a limited molecular genetic understanding of these bacteria relative to well-studied species such as *E. coli*. To increase our knowledge of SRB biology, we applied a high-throughput genetics driven approach, RB-TnSeq, to systematically identify mutant phenotypes for the commonly studied SRB *D. vulgaris* Hildenborough. From these phenotypes and comparative genomic analyses, we were able to make a number of new insights into the physiology and metabolism of DvH.

The large-scale genetic dataset we present for DvH can serve as a powerful tool for developing hypotheses regarding the functions of genes in this bacterium. Importantly, our dataset complements other resources available for this strain including gene regulatory and metabolic models. The DvH RB-TnSeq library in JW710 is readily available for other groups to perform additional genome-wide assays; furthermore the availability of archived single mutants (both targeted gene deletions and transposon insertion strains) can greatly accelerate follow-up studies on genes of interest.

## MATERIALS AND METHODS

### Bacterial strains and materials

The bacterial strains and oligonucleotides used in this study are listed in Supplementary Tables 4 and 5, respectively. Oligonucleotides for cloning were ordered from Life Technologies (www.thermofisher.com) and oligonucleotides for next-generation sequencing libraries preparation were ordered from Integrated DNA Technologies (www.IDT.com). We used the GoTaq® Green Master Mix (Promega) for colony PCRs; and Q5 hot start DNA polymerase (New England Biolabs) for all other PCR reactions. DNA fragments were purified with the QIAquick PCR purification or Gel extraction kit (Qiagen). T4 DNA ligase and buffer were purchased from NEB. Plasmid and genomic DNA isolations were carried out with the QIAprep Spin Miniprep Kit and the DNeasy Blood & Tissue Kit (Qiagen), respectively.

The DvH genome annotation used in this study includes protein-coding genes that are not in the current version in GenBank (GCF_000195755.1). These additional genes were identified by transcriptomics and proteomics evidence (Price et al., 2011), and each starts with the systematic name “DORF”. These annotations are included in the DvH information at MicrobesOnline (Dehal et al., 2010).

The genes neighborhood comparison shown in this study was achieved using the BioCyc Database collection and the Ensembl Bacteria browser (Howe et al., 2020; Karp et al., 2019).

### Growth conditions

*E. coli* conjugation strain APA766 was cultured in Luria-Bertani (LB) medium at 37°C supplemented with 100 μg/ml kanamycin and diaminopimelic acid (DAP) added to a final concentration of 300 μM. *D. vulgaris* Hildenborough strains were grown within an anaerobic chamber (Coy Laboratory Products, Grass Lake, MI) with an atmosphere of about 2% H_2_, 5% CO_2_, and 93% N_2_. DvH was grown in MO medium (Zane et al., 2010). When 60mM lactate (carbon source), 30mM sulfate (electron acceptor) and yeast extract (1 g/liter) were added, the medium was designated MOYLS4 medium. A description of the media used in this study and their composition is given in Supplementary Table 6. Other carbon sources and electron acceptors were used and their concentrations are detailed in Supplementary Table 3. All media were autoclaved, moved to the anaerobic chamber before cooling and amended with sodium sulfide (1mM) as reductant prior to inoculation. MOYLS4 agar plates were poured inside the anaerobic chamber 1 − 2 days prior to use. Culture growth was measured with a Bioscreen C instrument (Growth Curves USA) housed within the anaerobic chamber at 30°C.

### Transposon mutant library construction

We constructed *Tn*5 transposon mutant libraries in the wild-type and JW710 backgrounds with minor modifications to those previously described protocols (Wetmore et al., 2015). Our *Tn*5 transposon was a barcoded derivative of the *Tn*5-RL27 transposon (Larsen et al., 2002), which is itself a derivative of the *Tn*5 transposon (Reznikoff, 2008). Briefly, we conjugated mid-log-phase grown *E. coli* APA766 and DvH cells at 2:1 ratio on a 0.45 uM nylon membrane filter (Supor) overlaid on MOYLS4 agar plates. After 4 hours of anaerobic incubation (30°C), the filters were transferred into liquid MOYLS4 medium. After 4 hours of recovery at 30°C, the cells were then transferred into the same medium supplemented with G418 (400 ug/mL), and grown to saturation to select for G418-resistant transposon mutants. We made multiple, single-use glycerol stocks of the library and extracted genomic DNA for Tn-seq analysis. To map the genomic location of the transposon insertions and to link these insertions to their associated DNA barcode, we used the same Tn-seq protocol that was previously described (Wetmore et al., 2015).

### Identification of essential genes

Genes that are essential (or nearly so) for viability under the conditions used to select the mutants were identified with previously described criteria (Price et al., 2018; Rubin et al., 2015). Only protein-coding genes with at least 100 nucleotides of non-repetitive sequence (so that transposon insertions could be unambiguously mapped) were considered. A gene was deemed essential if the normalized density of insertions (dens, scaled so that the median is 1) and normalized reads per kilobase (normreads, scaled so that the median is 1 and also normalized for GC content) were both below 0.2 (see Supplementary Table 1), and if the mutants in that gene did not have sufficient abundance in Time0 samples to calculate fitness scores. As the wild-type strain was not used for fitness assays, we used the Time0 samples from the JW710 mutant library for this parameter in the wild-type essentiality calls. Of the 19 genes called as essential in the wild-type background but not in JW710, 18 were considered “nearly essential” because either dens or normreads were below 0.2 (Supplementary Table 1). From the Fels *et al*. TnSeq data (Fels et al., 2013), we determined the number of different positions with a central insertion (10 − 90% of gene length; we ignored insertion positions with only 1 read) and the total number of sequencing reads for each gene at these positions (Supplementary Table 1). Genes with 10 or more insertion positions and 100 or more total reads were considered to be dispensable for viability. To compare the 380 genes that were identified as likely essential in both the wild-type and JW710 backgrounds to essential genes in other bacteria, we used the Database of Essential Genes (Zhang et al., 2004), which contains lists of essential genes from dozens of bacteria. Specifically, we downloaded the latest version of DEG (version 15.2) and used BLASTp to search for homologs of the DVH proteins. We considered the DvH protein to have a homolog if percent identity to the DEG protein was at least 30%, and if the alignment covered at least 75% of the protein’s length in both the DvH and DEG proteins.

### Sequence differences between wild-type DvH and JW710

The genome of JW710 was sequenced on an Illumina MiSeq by the University of Missouri DNA Core. Within the Geneious software (v 8.1.9) and with default parameters, raw sequences from JW710 were trimmed to remove adapter sequences, mapped to the DvH genome reference sequence (NCBI Accession No. NC_002937.3 and NC_005863.1) with Bowtie2 (Langmead and Salzberg, 2012), and sequence variants were identified. To identify sequence variants in the wild-type DvH strain used in this study, we aligned the Tn-seq reads from the wild-type background (after removing the sequences corresponding to the transposon) against the published reference genome, and identified sequence variants as described above for JW710. This analysis covered about 75% of the genome.

### Genome-wide mutant fitness assays

We performed pooled mutant fitness assays as described previously (Price et al., 2018; Wetmore et al., 2015). Briefly, an aliquot of the full transposon mutant library was inoculated in MOYLS4 medium supplemented with G418 (400 ug/mL) and culture was left to grow anaerobically at 30°C until the cells reached mid-log phase. Samples of the culture of pooled mutants were collected as the “Time0” controls. The remaining culture was pelleted, washed twice with phosphate buffer, and finally resuspended in phosphate buffer (or in MOLS4 when formate-acetate was used as only carbon source). We inoculated the mutant pool in the selective medium at a starting optical density at 600 nm of 0.02. The fitness assays were grown in either 24-well microplates or in 15 mL tubes; all plates and tubes were equilibrated inside the anaerobic chamber prior use. After 4 to 6 population doublings of the mutant library, we collected “condition” samples. We then extracted genomic DNA from the Time0 and condition samples. The DNA barcodes were amplified and sequenced (BarSeq) with previously established protocols (Price et al., 2018; Wetmore et al., 2015).

### Data analysis

We calculated gene fitness scores as described (Wetmore et al., 2015). Briefly, strain fitness scores are calculated as the normalized log2 ratio of the abundance of the barcode after selection (condition) versus before (Time0). Gene fitness is computed as the weighted average of the fitness of the individual mutants; then, to correct for variation in copy number along the chromosome in growing cells, gene fitness values are normalized so that the running median along the main chromosome is zero; finally, the gene fitness values are normalized so that the mode (for genes on the chromosome) is zero. For each gene fitness score, we calculate a *t*-like test statistic to determine the significance of the measurement (Wetmore et al., 2015). For calculating the number of DvH genes with a significant phenotype in the entire dataset, we required |fitness| > 0.5 and |*t*| > 4. This analysis identified 1,137 genes with a phenotype in at least one of the 757 experiments. To determine the false discovery rate (FDR), we used comparisons among the Time0 samples, which are not expected to result in significant phenotypes. Among 83 Time0 comparisons, we identified 4 genes with a significant phenotype with the same thresholds for fitness and *t*. After correcting for having 757 condition experiments and only 83 Time0 samples, we estimated the FDR for genes with a significant phenotype as 3% (36/1,137).

DvH genes with a specific phenotype in an experiment were defined as: |fitness| > 1 and |*t*| > 5; |fitness| < 1 in at least 95% of experiments; and the fitness value in this experiment was more pronounced that most of its other fitness values (|fitness| > 95th percentile(|fitness|) + 0.5) (Price et al., 2018). Cofitness was calculated as the Pearson correlation coefficient between all 757 fitness measurements for each pair of genes (Price et al., 2018).

To infer the functions of DvH genes based on mutant phenotypes, we primarily used the fitness browser (fit.genomics.lbl.gov), which contains genome-wide mutant fitness data for 38 different bacteria and a number of interactive tools for data exploration (Price et al., 2018). To determine the current state of knowledge on individual DvH proteins and their homologs, we used PaperBLAST (http://papers.genomics.lbl.gov/) (Price and Arkin, 2017). We used TIGRFAMs to assign DvH proteins to different functional categories (primarily using main role categories) (Haft et al., 2013). RegPrecise (Novichkov et al., 2013) was accessed using MicrobesOnline or from github (https://github.com/SMRUCC/RegPrecise/tree/master/genomes).

To identify protein-coding genes with hypothetical or vague annotations, we matched their descriptions against text patterns such as “hypothetical”, “family”, or “membrane protein,” as described previously (Price et al., 2018).

## Data and software availability

The data and analyses described in this work can be accessed from different sources. First, the gene fitness values and their comparison to fitness data from other bacteria are available through the fitness browser (fit.genomics.lbl.gov, archived at https://doi.org/10.6084/m9.figshare.13172087.v1). This is the best location to access the data to examine the phenotypes of specific genes of interest or to BLAST a protein of interest against the entire dataset. Second, the gene fitness values, *t* scores, detailed metadata for all experiments, barcode counts, tables of genes with specific phenotypes and cofitness, and a single R image with all analyzed fitness data, are available at figshare (https://doi.org/10.6084/m9.figshare.13010285). Third, all of the TnSeq data is available from NCBI’s sequence read archive under project PRJNA666215. Fourth, genome sequencing data for strain JW710 is available at accession SRX9297596.

The software we used for analyzing the Tn-seq and BarSeq fitness data is available at https://bitbucket.org/berkeleylab/feba/.

## ACKNOWLEDGEMENTS

We thank Lara Rajeev for her help with anaerobic growth monitoring and Hans K. Carlson for helpful discussions and critical reading of the manuscript.

