## Supplementary Figures for "Large-scale Genetic Characterization of a Model Sulfate-Reducing Bacterium"

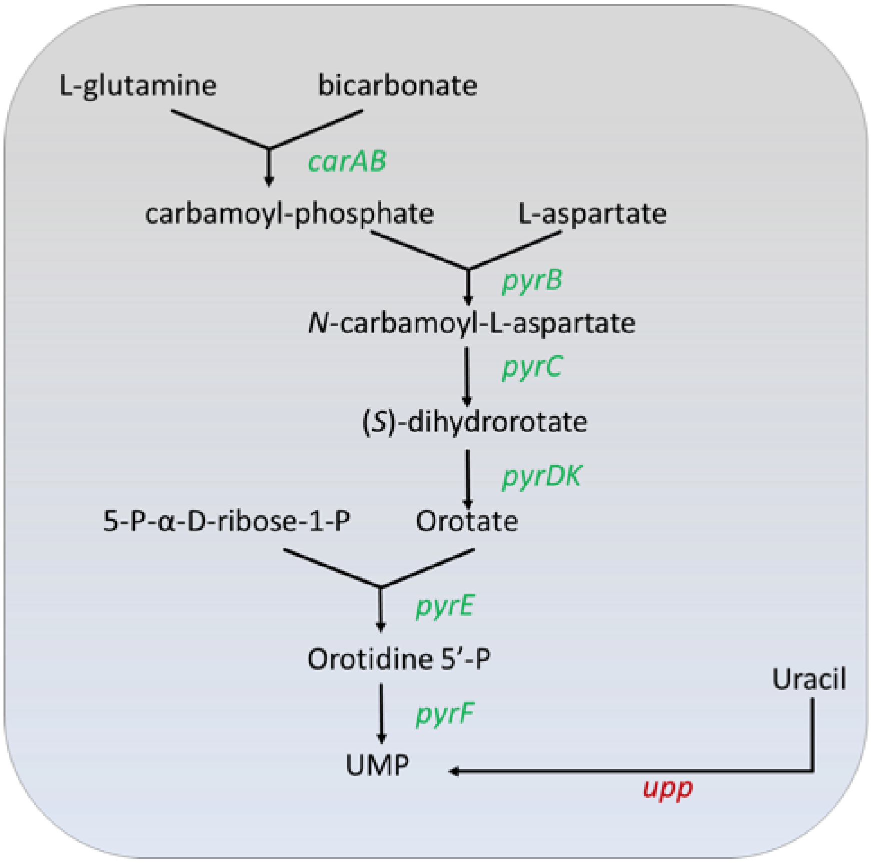


**Supplementary Figure 1.** Genes of the *de novo* UMP biosynthesis pathway are marked in green, all of these genes are essential for viability in the JW710 background that contains a deletion of *upp*. In contrast, all 8 of these genes are non-essential in the wild-type DvH background, as the intact *upp* provides a route to synthesize UMP.

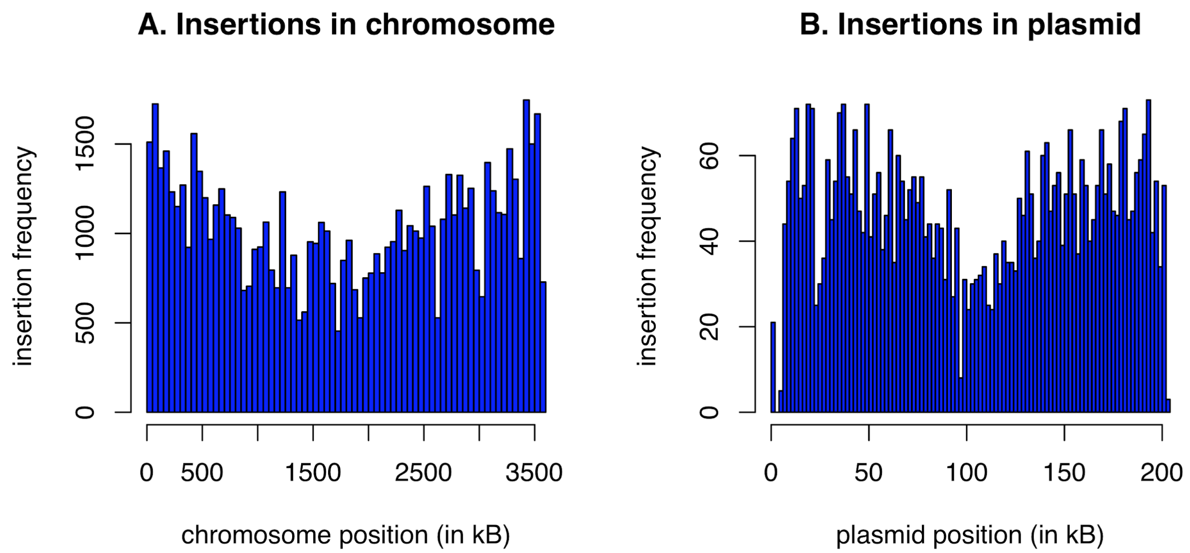


**Supplementary Figure 2.** Insertion coverage of mapped transposon insertions in the DvH JW710 RB-TnSeq mutant library across the chromosome (A) and megaplasmid (B).

| | Order | Family | Genus | Organisms |
|---|---|---|---|---|
| *Deltaproteobacteria* | *Desulfovibrionales* | *Desulfovibrionaceae* | *Desulfovibrio* | *Desulfovibrio vulgaris* DP4<br>*Desulfovibrio vulgaris* Hildenborough<br>*Desulfovibrio vulgaris* Miyazaki<br>*Desulfovibrio vulgaris* RCH1 |
| | | | | *Desulfovibrio alaskensis* G20 |
| | | | | *Desulfovibrio* sp. A2 |
| | | | | *Desulfovibrio piger* ATCC 29098 |
| | | | *Bilophila* | *Bilophila wadsworthia* ATCC 49260<br>*Bilophila wadsworthia* 3_1_6 |
| | | | | *Bilophila* sp.4_1_30 |
| | | | *Halodesulfovibrio* | *Halodesulfovibrio aestuarii* DSM 10141 |
| | | | | *Halodesulfovibrio marinisediminis* DSM 17456 |
| | | | | *Halodesulfovibrio spirochaetisodalis* |
| | | *Desulfohalobiaceae* | *Desulfovermiculus* | *Desulfovermiculus halophilus* DSM 18834 |
| | | | *Desulfothermus* | *Desulfothermus okinamensis* |
| | | *Desulfomicrobiaceae* | *Desulfomicrobium* | *Desulfomicrobium baculatum* DSM 4028 |
| | | | | *Desulfomicrobium escambiense* DSM 10707 |
| | | | | *Desulfomicrobium orale* DSM 12838 |
| | *Desulfobacterales* | *Desulfobacteraceae* | *Desulfobacter* | *Desulfobacter vibrioformis* DSM 8776 |
| | | | *Desulfobacterium* | *Desulfobacterium vacuolatum* DSM 3385 |
| | | | Desulfona | *Desulfona spongiiphila* |
| | | | *Candidatus magnetomorum* sp. HK1 | |
| | *Syntrophobacterales* | *Syntrophaceae* | *Syntrophus* | *Syntrophus aciditrophilus* SB<br>*Syntrophus gentianae* DSM 8423 |
| | | *Syntrophobacteraceae* | *Desulfoglaeba* | *Desulfoglaeba alkenexedens* ALDC |
| | | | *Syntrophobacter* | *Syntrophobacter fumaroxidans* MPOB |
| *Chrysiogenetes* | *Chrysiogenales* | *Chrysiogenaceae* | *Desulfurisprillum* | *Desulfurisprillum indicum* S5 |

Biotin gene cluster

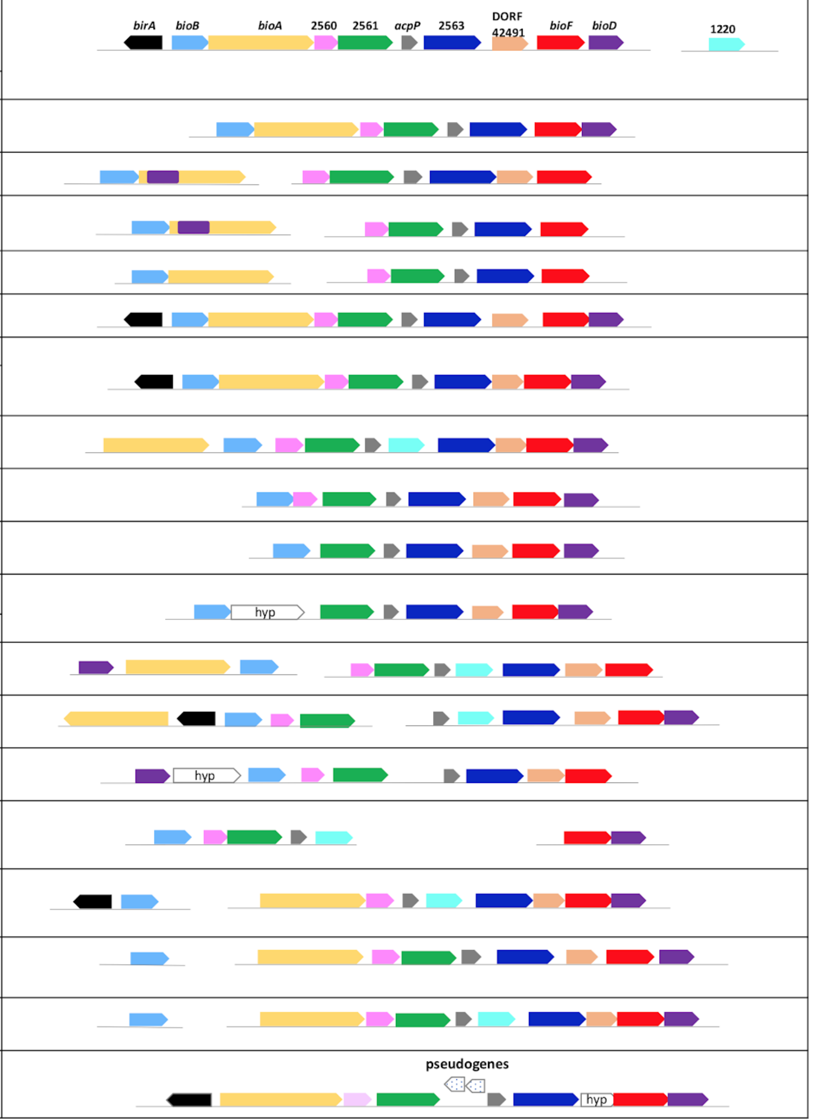

**Supplementary Figure 3.** Biotin cluster genes conservation in Deltaproteobacteria and Chrysiogenetes. The *birA* genes are shown only when located in the direct vicinity of a *bio* gene. In *Halodesulfovibrio spirochaetisodalis* genome, the gene homolog to DvH DORF41491 was added to the original annotation. The regions of *Desulfobacterium vacuolatum* DSM 3385 genome shown in the figure represent the end and beginning of two separate contigs.

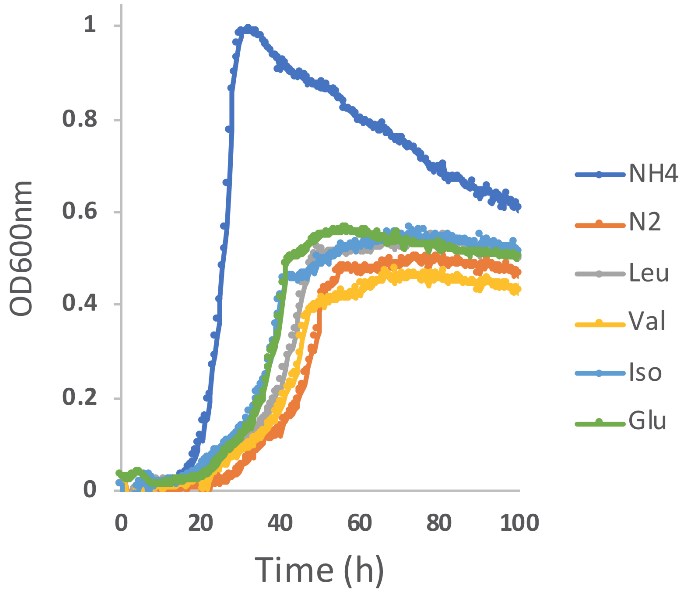


**Supplementary Figure 4.** Growth of DvH in lactate-sulfate minimal medium with different compounds as the sole source of nitrogen: $N_2$ (90%atm), $NH_4$, valine, leucine, isoleucine and glutamate (20mM). Measurements were made in a Bioscreen growth analysis system and each curve is the average of four replicates.
